# Role of SARS-CoV-2 in altering the RNA binding protein and miRNA directed post-transcriptional regulatory networks in humans

**DOI:** 10.1101/2020.07.06.190348

**Authors:** Rajneesh Srivastava, Swapna Vidhur Daulatabad, Mansi Srivastava, Sarath Chandra Janga

## Abstract

The outbreak of a novel coronavirus SARS-CoV-2 responsible for COVID-19 pandemic has caused worldwide public health emergency. Due to the constantly evolving nature of the coronaviruses, SARS-CoV-2 mediated alteration on post-transcriptional gene regulation across human tissues remains elusive. In this study, we analyze publicly available genomic datasets to systematically dissect the crosstalk and dysregulation of human post-transcriptional regulatory networks governed by RNA binding proteins (RBPs) and micro-RNAs (miRs), due to SARS-CoV-2 infection. We uncovered that 13 out of 29 SARS-CoV-2 encoded proteins directly interact with 51 human RBPs of which majority of them were abundantly expressed in gonadal tissues and immune cells. We further performed a functional analysis of differentially expressed genes in mock-treated versus SARS-CoV-2 infected lung cells that revealed enrichment for immune response, cytokine-mediated signaling, and metabolism associated genes. This study also characterized the alternative splicing events in SARS-CoV-2 infected cells compared to control demonstrating that skipped exons and mutually exclusive exons were the most abundant events that potentially contributed to differential outcomes in response to viral infection. Motif enrichment analysis on the RNA genomic sequence of SARS-CoV-2 clearly revealed the enrichment for RBPs such as SRSFs, PCBPs, ELAVs, and HNRNPs suggesting the sponging of RBPs by SARS-CoV-2 genome. A similar analysis to study the interactions of miRs with SARS-CoV-2 revealed functionally important miRs that were highly expressed in immune cells, suggesting that these interactions may contribute to the progression of the viral infection and modulate host immune response across other human tissues. Given the need to understand the interactions of SARS-CoV-2 with key post-transcriptional regulators in the human genome, this study provides a systematic computational analysis to dissect the role of dysregulated post-transcriptional regulatory networks controlled by RBPs and miRs, across tissues types during SARS-CoV-2 infection.

## 1. Introduction

An outbreak of coronavirus disease (COVID-19) caused by the newly discovered severe acute respiratory syndrome coronavirus (SARS-CoV-2), started in December 2019, in the city of Wuhan, Hubei province, China. As of September 16, 2020, COVID-19 has expanded globally with more than 30 million confirmed cases with over 944,000 deaths worldwide, imposing an unprecedented threat to public health (https://www.worldometers.info/coronavirus/). In the past two decades, coronavirus outbreak has resulted in viral epidemics including severe acute respiratory syndrome (SARS-CoV) in 2002 with fatality of 10% and middle east respiratory syndrome (MERS-CoV) in 2012 with fatality of 36% [1–4]. Both SARS-CoV and MERS-CoV were zoonotic viruses originating in bat and camels respectively [5, 6]. However, the recurring emergence of highly pathogenic SARS-CoV, MERS-CoV and now SARS-CoV-2 have indicated the potential for cross-species transmission of these viruses thus raising a serious public health concern [7, 8]. SARS CoV-2 shares a sequence similarity of 80% and 50% with previously identified SARS-CoV and MERS-CoV respectively [9–12]. Since its emergence, rapid efforts have illustrated the molecular features of SARS-CoV-2 that enables it to hijack the host cellular machinery and facilitates its genomic replication and assembly into new virions during the infection process [13–16].

Coronavirus carries the largest genome among all RNA viruses, ranging from 26 to 32 kilobases in length [12]. This virus has a characteristic “crown” like appearance under two-dimensional transmission electron microscopy. SARS-CoV-2 is an enveloped positive-sense, single-stranded ribonucleic acid (RNA) coronavirus that belongs to the genus beta-coronavirus. Upon entry in the cell, SARS-CoV-2 RNA is translated into non-structural proteins (nsps) from two open reading frames (ORFs), ORF1a, and ORF1b [17, 18]. The ORF1a produces polypeptide 1a that is cleaved further into 11 nsps, while ORF1b, yields polypeptide 1ab, that is cleaved into 16 nsps [17, 18]. Since SARS-CoV-2 utilizes human machinery to translate its RNA after the entry into the cell, it could possibly impact several RNA-binding proteins from the host to bind the viral genome resulting in altered post-transcriptional regulation. Next, the viral genome is used as the template for replication and transcription, mediated by non-structural protein, RNA-dependent RNA polymerase (RdRP) [18, 19]. SARS-CoV-2 encodes four main structural proteins: spike (S), envelope (E), membrane (M), and nucleocapsid (N) that are conserved and several other accessory proteins (3a, 6, 7a, 7b, 8, and 10) according to the current annotation (GenBank: NC_045512.2) [17, 20]. The spike protein, that has evolved the most during the COVID-19 outbreak, enables the virus to bind to angiotensin-converting enzyme 2 (ACE2) on the host cell membrane, following which it undergoes structural changes and subsequently allows the viral genome to make its way inside the host cell [21]. Infections caused by these viruses result in severe pneumonia, fever and breathing difficulty [22].

Protein-protein interaction map between SARS-CoV-2 and human proteins published recently has revealed several important targets for drug repurposing [23]. Given the evolving nature of coronaviruses that results in frequent genetic diversity in their genome, it is crucial to identify the regulators in humans that interact with the viral genome and their cross talk that results in altered regulatory mechanisms in the host during the infection process. Therefore, it is imperative to investigate the interacting post-transcriptional regulators that asset these viral proteins in different tissues.

RNA-binding proteins (RBPs) are a class of proteins in humans that bind to single or double stranded RNA and facilitate the formation of ribonucleoprotein complexes [24–26]. In addition to RBPs, micro-RNAs (miRs) that belong to a class of non-coding RNAs also interact with target RNAs to regulate cognate RNA expression [27, 28]. Both RBPs and miRs have been widely recognized in regulating the post-transcriptional gene regulatory network in humans [29–31]. Dysregulated RBPs and miRs have been shown to contribute significantly to altered regulatory network in a plethora of diseases such as cancer, genetic diseases and viral infections [32–38]. Previous studies have shown that human RBPs including heterogeneous Nuclear Ribonucleoprotein family (hnRNPA1 and hnRNPAQ), polypyrimidine tract-binding protein (PTB), Serine/Arginine-Rich Splicing Factor 7 (SRSF7) and Transformer 2 Alpha Homolog (TRA2A) interact with coronavirus RNA [39–44]. Likewise, other reports have demonstrated the potential interaction between human miRNA and viral genome including variety of coronaviruses [41, 45, 46]. However, the potential RBPs and miRs that interact with SARS-CoV-2 and their implication in viral pathogenesis has been poorly understood.

Currently, there are no proven anti-viral therapeutics that are effective against the novel coronavirus. Although, analysis of therapeutic targets for SARS-CoV-2 has been conducted to identify potential drugs by computational methods [47], the targets have not clinically approved for therapeutic applications. Alternative therapeutics like angiotensin receptor blockers have been identified as tentative target candidates but have shown concerns associated with the loss of angiotensin functions crucial for cell [48]. Therefore, to devise effective therapeutics, there is a need to determine the cellular targets in humans that interact with the virus and result in altered functional outcomes. In this study, we uncovered that several human RBPs and miRNAs harbor abundant binding sites across the SARS-CoV-2 genome, illustrating the titration of post-transcriptional regulators. Interestingly, we show that most of these regulators were predominantly expressed in gonadal tissues, adrenal, pancreas, and blood cells. Overall, this study will bridge the gap in our understanding of the impact of SARS-CoV-2 infection on post-transcriptional regulatory networks.

## 2. Results and discussion

### 2.1. Protein-protein interaction network analysis for SARS-CoV-2 viral proteins reveals an extensive and direct set of interactions for scores of functionally important human RBPs

We obtained the affinity purification-mass spectrometry (AP-MS) based SARS-CoV-2 and human proteins interaction network established in HEK293 cells [23] and investigated the human RBPs that directly interact with viral proteins. Our analysis revealed that SARS-CoV-2 encoded proteins interact directly with 51 human RBPs (Fig. 1A). We observed that these primary interacting RBPs were proven to serve several vital functions in the cells such as polyadenylate binding protein 4 (PABP-4) and Dead-box RNA helicases (DDX21 and DDX10), enzymes involved in translation machinery such as eukaryotic translation initiation factor 4H (EIF4H) and ribosomal protein L36 (RPL36) (Fig. 1A). Among the direct interactors, the highly abundant cytoplasmic PABPs, known to bind the 3’ polyA tail on eukaryotic mRNAs, has previously been reported to interact with polyA tail in bovine coronavirus and mouse hepatitis virus [49–51]. Since SARS-CoV-2 is also composed of polyadenylated RNA, it is likely that the host PABP could modulate the translation of coronavirus genome through polyA binding. DDX10, another primary interactor observed in the analyzed dataset, has been reported to interact with SARS-CoV-2 non-structural protein 8 (nsp8) [52] suggesting that the identified host RBPs could be implicated in the regulatory processes of SARS-CoV-2 genome synthesis. EIF4H, also found as one of the primary interactors, was reported to interact with SARS-CoV-2 non-structural protein 9 (nsp9) in a recently published study [23]. Furthermore, among the immediate interactions, we also found human RBPs such as signal recognition particle 19 (SRP19 and SRP54) and Golgin subfamily B member 1 (GOLGB1) that have been well recognized for co-translational protein targeting to membrane and endoplasmic reticulum to golgi vesicle-mediated transport [53, 54] (Fig. 1A). These results suggest that several human RBPs that come in direct contact with SARS-CoV-2 proteins could contribute to virus assembly and export and could therefore be implicated as therapeutic targets. However, such findings require in-depth experimental validation in a tissue-specific context to support the functional involvement of the identified RBPs in response to SARS-CoV-2 infection.

**Figure 1.**
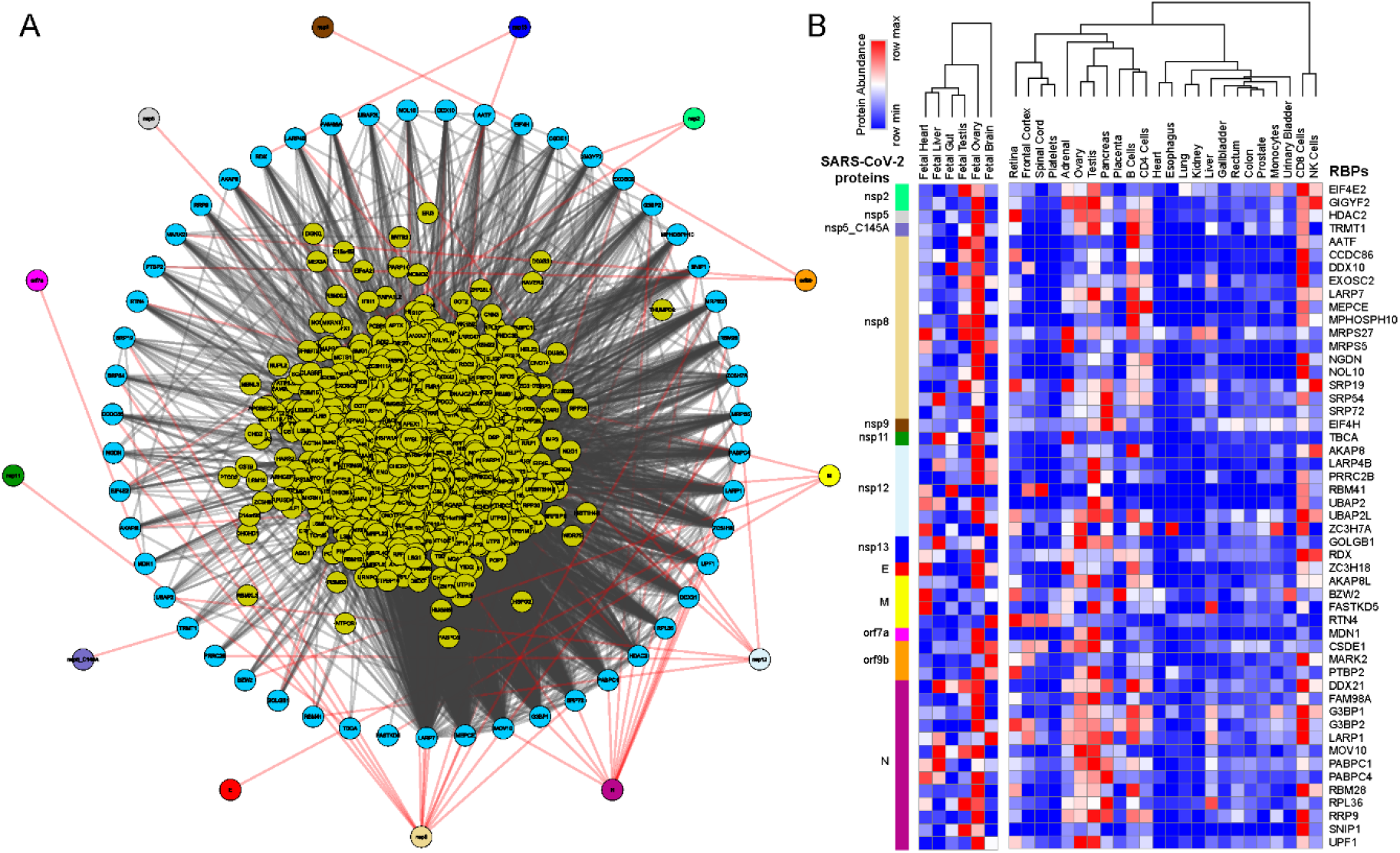
Protein-protein interaction network analysis suggest an immediate interaction of human RBPs with SARS-CoV-2 viral proteins (A) An integrated SARS-CoV-2 – human RBP interaction network. We obtained the MS-based SARS-CoV-2 viral protein to human protein interaction network established in HEK293 cells and integrated with 1st neighbor interacting RBPs (obtained from BioGRID). (B) Protein abundance of SARS-CoV-2 interacting RBPs across human tissues. Expression data were obtained from human protein map and row normalized. SARS-CoV-2 proteins were color coded and highlighted in the network.

Further, we observed that among the SARS-CoV-2 encoded proteins, majority of the direct interactions occurred with a range of non-structural proteins (nsp 2,5,8,9,11,13) that contribute to viral replication and transcription along with structural proteins (E,N and M) (Fig. 1A). Therefore, it is likely that the identified human RBPs that interact with viral proteins assist in the viral pathogenesis. Furthermore, we also observed that ~65% of annotated human RBPs [55] were in immediate neighborhood (obtained from BioGRID [56], shown at the center in Fig. 1A) of virus-protein interacting RBPs and could indirectly regulate the SARS-CoV-2 proteins. Overall, the comprehensive direct and indirect interactions between human RBPs and SARS-CoV-2 proteins is likely to rewire the post-transcriptional gene regulatory mechanisms in human cells.

Next, we examined the abundance of SARS-CoV-2 interacting RBPs across human tissues using the protein expression data from human proteome map [57]. Our results suggest that majority of the human RBPs that have direct interaction with SARS-CoV-2 proteins were predominantly expressed in gonadal tissues (Testis and Ovary) (Fig. 1B). These findings agreed with a recent study showing that male reproductive systems are vulnerable to SARS-CoV-2 infection, that was evident by dramatic changes in sex hormones of the infected patients suggesting gonadal function impairment [58, 59]. Additionally, we also found that, these SARS-CoV-2 interacting human RBPs were showing relatively higher expression in immune cell types such as T cells (CD4+ and CD8+) and NK cells that are a part of innate anti-viral immune response (Fig. 1B). Our observations are supported by recently published studies suggesting that T cells, CD4+ T cells, and CD8+ T cells play a significant antiviral role during SARS-CoV-2 infection [20, 60–63]. Overall, the results from this analysis provide a systematic dissection of potential RBPs in humans interacting with SARS-CoV-2 proteins across tissues.

### 2.2. SARS-CoV-2 infected lung epithelial cells exhibit gene expression alterations in several immunological and metabolic pathways

Due to rapidly evolving nature of SARS-CoV-2, the transcriptomic alterations contributed by the virus in human remains unclear. To gain insight into the effect of SARS-CoV-2 infection on host gene expression, we obtained the raw RNA sequencing data in normal versus SARS-CoV-2 infected human bronchial epithelial (NHBE) cells [64] deposited in Gene Expression Omnibus (GEO) [65]. We investigated the gene expression levels between the mock treated versus SARS-CoV-2 infected NHBE cells and identified 327 differentially expressed genes at 5% FDR (among which ~67% shows >1.25 fold elevated expression in SARS-CoV-2 infected NHBE cells as shown in Fig. S1). We conducted functional pathways associated with these differentially expressed genes (Fig. 2A, Fig. S1) and identified 327 differentially expressed genes (at 5% FDR) using ClueGO [66] revealed an enrichment for immune response, cytokine mediated signaling, inflammatory response and metabolism associated genes (Fig. 2A, Table S1).

**Figure 2.**
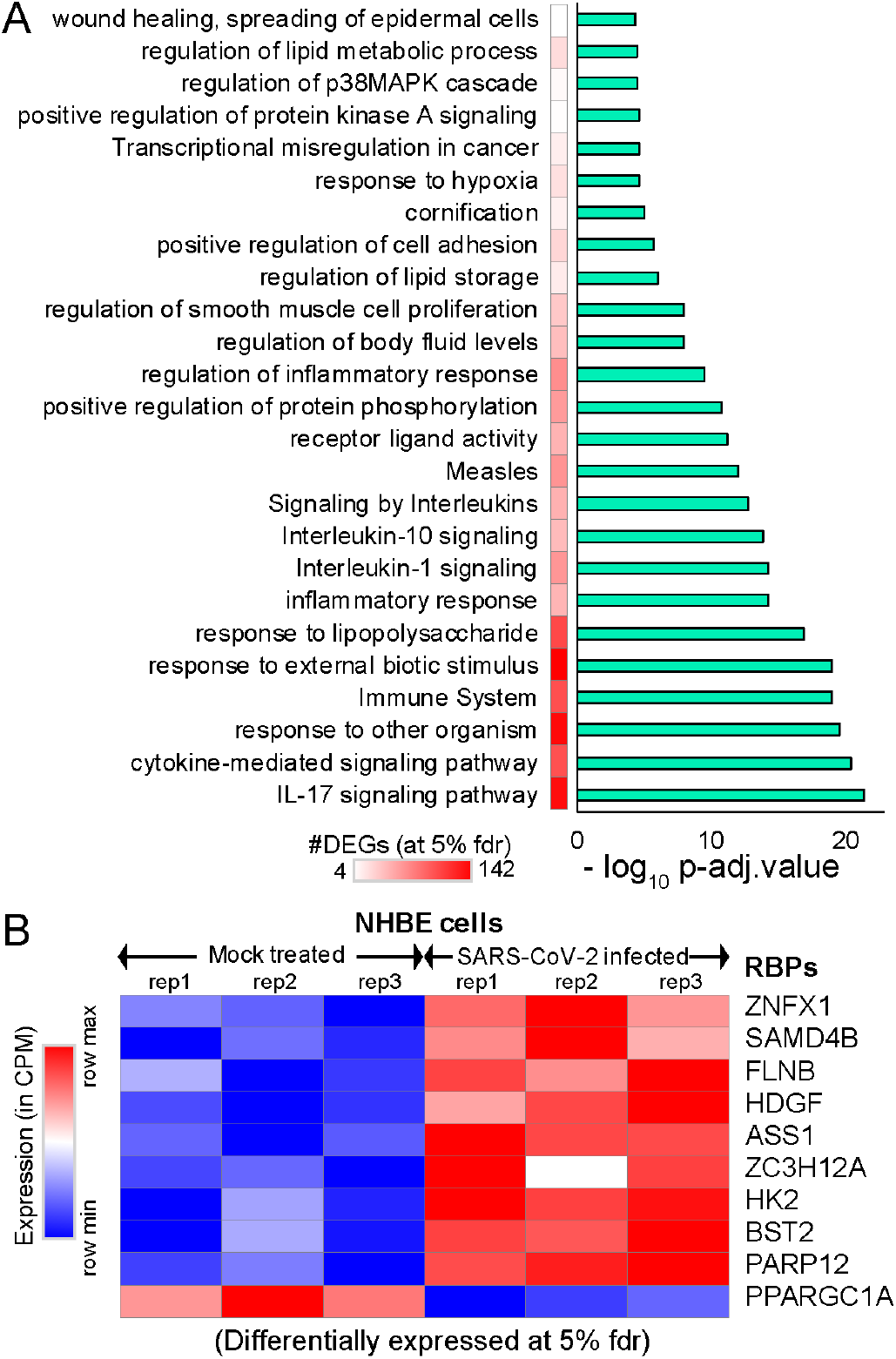
Differential expression analysis of mock treated versus SARS-CoV-2 infected primary human lung epithelial cells. (A) Bar-plot illustrating the significant pathways obtained from GO-term based functional grouping of differentially expressed genes (DEGs) at 5% fdr using ClueGO analysis (Cytoscape plugin) (B) Row normalized expression profile of differentially expressed RBPs in mock treated and SARS-CoV-2 infected primary human lung epithelial cells (in biological triplicates).

We observed a significant enrichment for the IL-17 pathway associated genes during SARS-CoV-2 infection (Fig. 2A). Our observation of overrepresented IL-17 pathway was in accordance with recent studies that show overactivation of IL-17 producing Th17 cells during severe immune injury in SARS-CoV-2 patients [67–69]. In addition to this, a recent review has summarized that targeting IL-17 is immunologically plausible as a therapeutic strategy to prevent acute respiratory distress syndrome (ARDS) during SARS-CoV-2 infection, based on previous evidence that mice lacking functional IL-17 receptor (Il17ra−/−) signaling were shown to be more susceptible than wild-type mice to secondary pneumonia following infection with influenza A [70, 71]. Thus, our analysis demonstrates the key role of several pathways including IL-17 cytokine response in SARS-CoV-2 infected cells.

Additional analysis also provided parallel support for the dysregulation of multiple RNA binding proteins (FLNB, HDGF, ASS1, ZC3H12A, HK2, BST2, PPARGC1A) involved in immune response, cytokine mediated signaling, and metabolism, when RNA-sequencing data from mock treated versus SARS-CoV-2 infected lung cells was employed (Fig. 2B). Dysregulated expression of multiple RBPs implicated in vital cellular functions suggests that the virus may hijack the host cellular machinery by modulating the expression of key RBPs. We also observed that six of these differentially expressed genes that encode for RBPs were involved in at least 30% of the overrepresented pathways. Overall, our results imply that differentially expressed RNA-binding proteins in SARS-CoV-2 infected cells may contribute to alterations in the post-transcriptional regulatory networks governed by them.

### 2.3. Alternative splicing analysis revealed an abundance of skipped and mutually exclusive exons in human transcripts during SARS-CoV-2 infection in lung epithelial cells

Alternative splicing is a principal mechanism that contributes to protein diversity in eukaryotes, while regulating physiologically important immune responses during bacterial and viral infections [72]. Viral infections have been shown to cause global changes in the alternative splicing signatures in the infected cells that may arise due to intrinsic factors like polymorphism at the splice sites or due to direct intervention by virulence factors [73–75]. A previous study on virus-host interactions has demonstrated that human coronavirus targets various signaling pathways of ER stress resulting in differential splicing outcomes [76]. Another study has shown that deletion of E protein in recombinant SARS-CoV resulted in significant XBP1 gene splicing and higher rate of apoptosis, suggesting that coronavirus infected cells are susceptible to differential splicing events [77]. Therefore, we next investigated the alternative splicing events in mock vs SARS-CoV-2 treated NHBE cells using rMATS (replicate Multivariate Analysis of Transcript Splicing) [78] (see Materials and Methods).

Our analysis revealed an abundance for skipped and mutually exclusive exonic events in the genes exhibiting alternative splicing events during SARS-CoV-2 infection at 5% fdr (Fig. 3A, Table S2). We also observed that 81 of the alternatively spliced genes encoded for RBPs (indicated in blue, Fig. 3A) and hence could result in altering the downstream post-transcriptional regulatory networks in SARS-CoV-2 infected cells (Fig. 3A).

**Figure 3.**
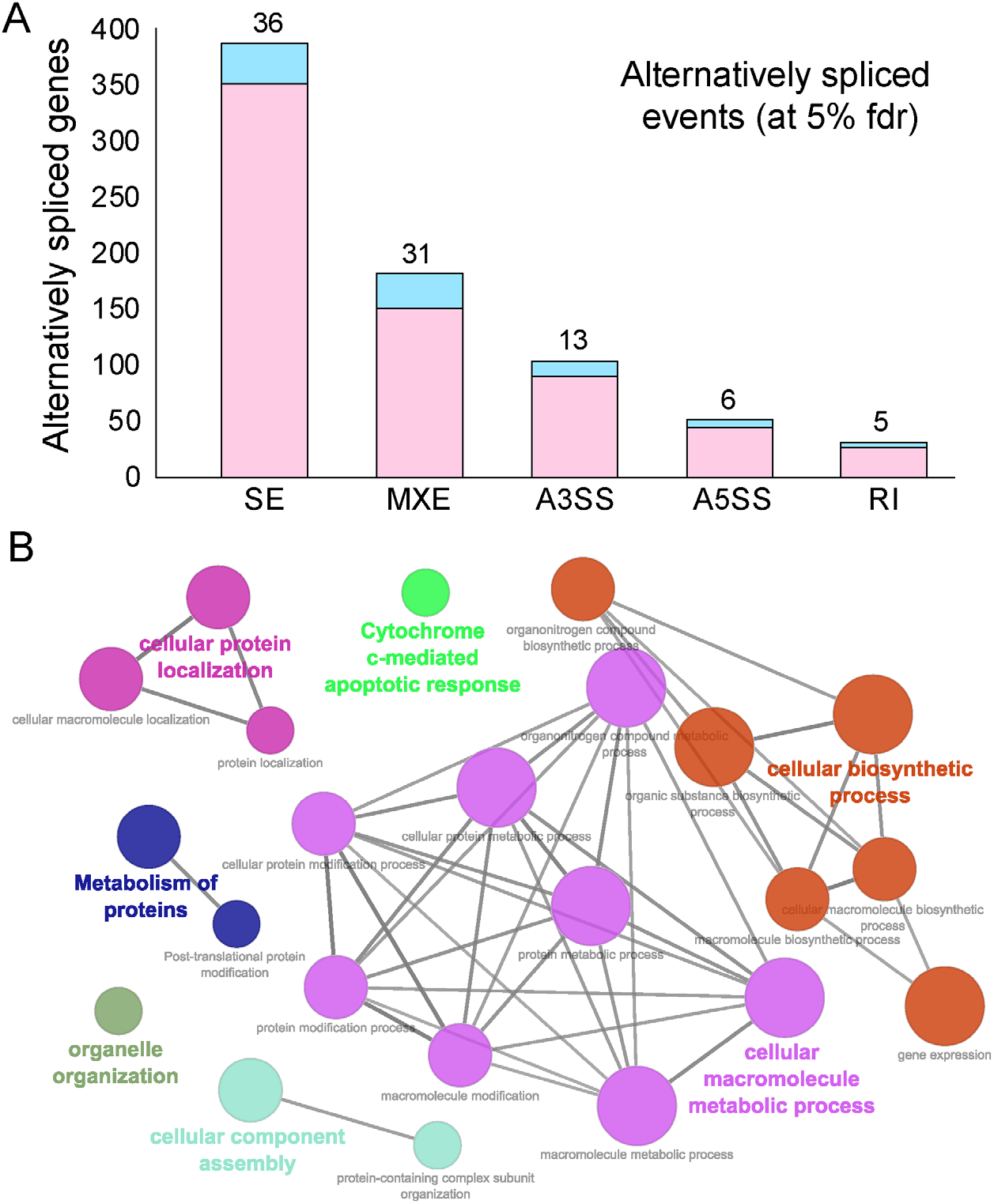
Alternative splicing events during SARS-CoV-2 infection. (A) Bar plot showing the genes (RBP encoding genes in blue) exhibiting alternative splicing during SARS-CoV-2 infection in primary human lung epithelial cells (at 5% fdr). (B) Clustered GO-term network obtained from function annotation analysis and grouping of the GO-term for the genes exhibiting alternative splicing using ClueGO (cytoscape plugin). Significant clustering (adj. p < 1e-05) of functional groups were color coded by functional annotation of the enriched GO-biological processes, with size of the nodes indicating the level of significant association of genes per GO-term, were shown.

These findings enhance our mechanistic understanding of the SARS-CoV-2 induced alternative splicing dysregulation in human cells and could be critical for developing novel therapeutic strategies. Next, we identified the functional annotation of the enriched GO-terms related to the genes exhibiting alternative splicing using ClueGO [66]. Our data revealed that majority of these genes were enriched for vital biological processes including cellular protein localization, protein metabolism, organelle organization, cellular biosynthetic process, cellular component assembly and cytochrome c-mediated apoptotic response (Fig. 3B, Table S3). These findings suggest that processes that contribute to cell structure, interaction, and growth upon SARS-CoV-2 infection via alternatively spliced genes could contribute to rewiring of post-transcriptional network. In summary, this analysis provides a clustered network of enriched biological functions that could be significantly dysregulated in SARS-CoV-2 infected cells

### 2.4. Motif enrichment analysis reveals potential human RBPs titrated by SARS-CoV-2 viral genome

SARS-CoV-2 genome is the largest among the coronavirus family (~30kb) and has been attributed to enhanced virus pathogenicity in the newly evolved strains of COVID-19 pandemic [18, 79]. Among the host derived cellular factors, RBPs have been recognized as active participants in all steps of viral infection [34, 80, 81]. A recent review has shown linkage of 472 human proteins with viruses through unconventional RNA binding domains [81]. However, the role of RNA binding proteins during viral pathogenesis has remained an unappreciated domain in viral research. Therefore, in the present research, we conducted a systematic and comprehensive bioinformatic study to investigate the RBPs that could potentially bind on RNA genome of SARS-CoV-2 by motif enrichment analysis of human RBPs using FIMO [82] (see Materials and Methods). Motif analysis for RBPs with established position specific weight matrices (PWMs) revealed significant number of binding sites spread across the SARS-CoV-2 genome illustrating the possible titration of post-transcriptional regulators by viral genome (Fig. 4A, Fig. S2, Table S4). To date, the PWMs are available only for a small fraction of the experimentally known human RBPs [83]. Thus, our analysis represents an understanding of post-transcriptional interactions for ~7.5% of total RBPs collected from multiple studies [24, 55, 84]. Importantly, the binding pattern of the RBP motifs across the entire normalized length of the virus suggests that, several of human RBPs could be titrated across the viral genome (Fig. 4A, Table S4). Our results showed an enrichment for RBPs such as SRSFs, PCBPs, ELAVs and HNRNPs being most likely to get sponged on the viral genome (Fig. 4B). Our observation that specific RBPs such as SRSF7, HNRNPA1 and TRA2A with well-known role in splicing exhibited binding sites on SARS-CoV-2 RNA, is in agreement with a recently published study [85]. We found that most of these RBPs were abundantly expressed in gonadal tissues, adrenal tissues, pancreas, and immune cells including B cells, CD4+ T cells and NK cells (Fig. 4B). Our analysis revealed that several members of SRSF family (SRSF 1,2,3,7,10) were potentially sponged by SARS-CoV-2 genome. Although SRSF2 has been reported to be predominantly nuclear, there is evidence for other SRSFs to be present in the cytoplasmic compartment by nuclear to cytoplasmic shuttling [86–91]. Therefore, such cytoplasmic binding and sequestration of SRSFs by SARS-CoV-2 RNA could contribute to dysregulation of host RNA targets. Furthermore, previous reports indicate that plus-stranded RNA viruses, such as polioviruses, utilize cellular factor PCBP for its translation and replication [44]. Therefore, we speculate that SARS-CoV-2 could also utilize host cell RBPs such as PCBPs to facilitate its genomic replication and translation. In addition to splicing factors and poly (rC) binding proteins, other cellular RBPs such as ELAV1 that regulates stability of host transcripts could also stabilize viral RNAs by sequestration on the viral genome [81, 92]. In summary, these findings suggest that SARS-CoV-2 could sponge human RBPs on its RNA resulting in altered post-transcriptional gene regulatory network in the host cells. Targeting host proteins has been appreciated as an effective strategy to combat a wide range of viral infections and therefore an understanding of the potential RBPs that are likely sponged by the viral genome is crucial to develop novel therapeutics [35].

**Figure 4.**
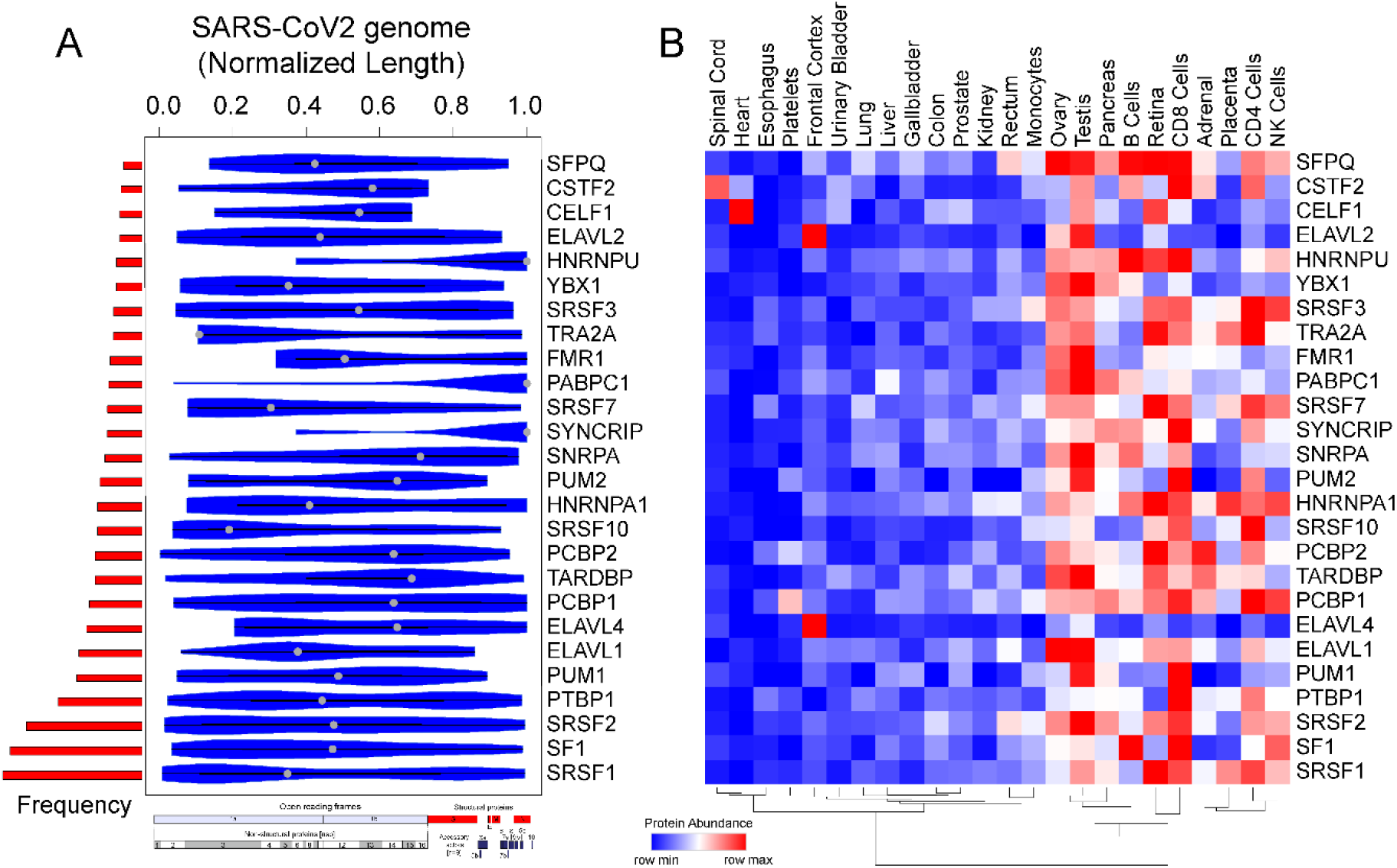
Motif enrichment analysis reveals potential human RBPs titrated by SARS-CoV-2 viral genome. (A) Violin plot shows the statistically significant (p-value < e-05) preferential binding profile of RBP motifs (sorted by frequency of binding and greater than 10 sites) across the SARS-CoV-2 viral genome (length normalized) identified using FIMO. (B) Hierarchically clustered heatmap showing the protein abundance (row normalized) of RBPs across tissues.

### 2.5. SARS-CoV-2 genome titrates the abundance of functionally important miRs in human tissue

Micro RNAs (miRs) are small non-coding RNA molecules that function as central regulators of post-transcriptional gene regulation. Human miRs have been associated with a variety of pathophysiological pathways and demonstrate differential expression during viral infections [93, 94]. Recently, a few computational studies have shed light on the interplay between human miRs and SARS-CoV-2 target genes indicating a crucial role in regulating the viral load in host cells [95, 96]. However, a comprehensive understanding of the functional role of host miRs during SARS-CoV-2 infection has remained elusive until now. Recently, a machine learning based study predicted that miRs could impact SARS-CoV-2 infection through several mechanisms such as interfering with replication, translation and by modulation of the host gene expression [95]. In this study, we used computational approach to investigate the potential binding sites of human miRs in SARS-CoV-2 genome using FIMO [82]. We identified 22 miRNAs that could potentially bind throughout the length of the SARS-CoV-2 viral genome (Fig. 5A, Table S5). Among the human miRs likely to be sequestered on SARS-CoV-2 genome, miR-374a-3p has recently been predicted to target SARS-CoV-2 gene, in particular it has been shown to target gene encoding the spike protein which is essential for virus entry into the host cell [95, 97]. In addition to targeting the spike protein encoding genes, miR-374a-3p is also predicted to target the ORF1ab in the SARS-CoV-2 genome that encodes for 5’viral replicase, based on the function similarity of SARS-CoV-2 coding genes with SARS-CoV [98]. Our data suggests that miRs can be bound by SARS-CoV-2 RNA, supporting a likely model where host miRs can be sponged by SARS-CoV-2 and thereby contributing to decreased binding of miRs to the human mRNA targets resulting in altering the expression patterns of human genes. Although less likely, it is also possible that that nuclear pre-processed miRNAs complimentary to SARS-CoV-2 RNA could be sequestered by viral RNA and because of this binding, miRNAs may not be able to do post-transcriptional gene silencing, thereby increasing the target gene expression. In either case, these results suggest that several important miRs are likely being titrated by SARS-CoV-2 genome that could result in dysregulation of post-transcriptional networks in the infected cells and could be attributed to viral pathogenesis. However, these results require experimental validation to conclude their role in SARS-CoV-2 infection.

**Figure 5.**
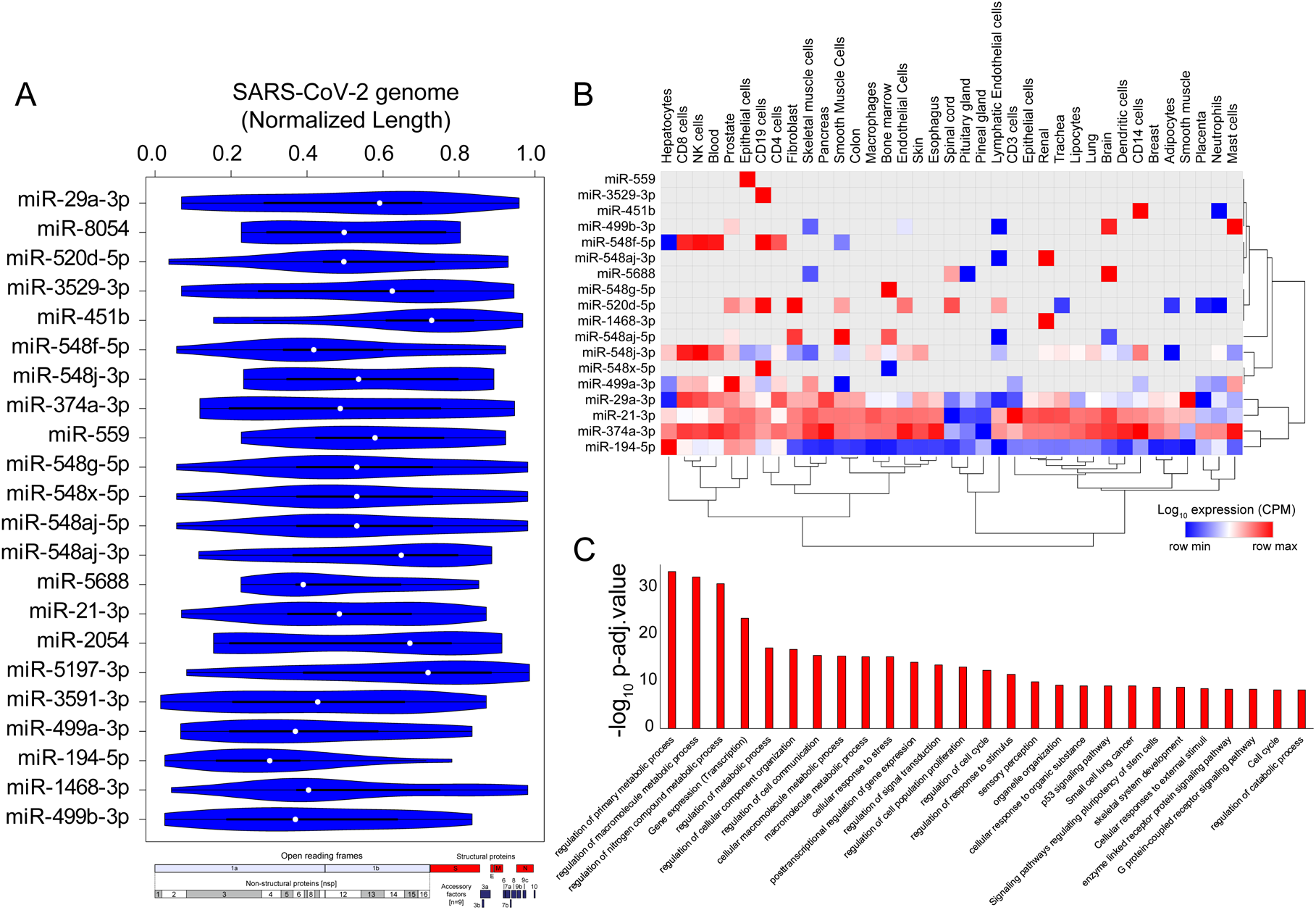
SARS-CoV-2 genome titrates the abundance of functionally important miRs in human tissue Violin plot shows the statistically significant (p-value < e-05) preferential binding profile of miR-motifs (sorted by frequency of binding >15 sites) across the SARS-CoV-2 viral genome (length normalized) identified using FIMO. (B) Hierarchically clustered heatmap showing the log10 expression (CPM, row normalized) of miRs across tissues. (C) Bar-plot illustrating the significant biological processes, obtained from gene ontology enrichment based functional grouping of miR target genes (obtained from miRNet). Significant clustering (adj. p < 1e-10) of genes enriched in GO-biological processes generated by ClueGO analysis (Cytoscape plugin).

Majority of the identified miRs were highly expressed in immune cells including CD8+T cells, CD4+T, NK cells, CD14 cells and mast cells, suggesting that these miRs might contribute to altered post-transcriptional regulation in specialized immune cells and assist in the progression of viral infection and host immune response across other human tissues (Fig. 5B). Our results also indicate that the high confident genes targeted by these sponged miRs were enriched for functional themes including ‘regulation of metabolic processes’, ‘post transcriptional gene regulation’ and ‘cell to cell communication’ suggestive of a large-scale dysregulation across tissues (Fig. 5C, Table S6). Overall, our results present a comprehensive analysis of the miRs being potentially titrated on the viral genome resulting in altered post-transcriptional gene regulation. These findings provide enhances our understanding of miR associated mechanism in SARS-CoV-2 pathogenesis and could provide important clues for designing RNA based therapeutics.

## 3. Conclusion

Our analysis integrates a comprehensive interaction network to map the immediate interactions between SARS-CoV-2 genome and proteome with human post-transcriptional regulators such as RBPs and miRNAs along with their tissue specific expression and functional annotations. Given the importance of developing effective therapeutic strategies in the current pandemic, understanding the effect of SARS-CoV-2 infection on human transcriptional and post-transcriptional regulatory networks is crucial for identifying effective drug targets. To delineate the impact of SARS-CoV-2 on host cells post-transcriptional gene regulatory network, we integrated the interactions between SARS-CoV-2 encoded proteins with human RNA binding proteins. Our findings indicate 51 human RBPs (including PABP-4, DDX21, DDX10, EIF4H) that interact directly with the viral structural and non-structural proteins, that in turn interact indirectly with ~65% other secondary neighbor RBPs. Thus, these findings suggest a comprehensive network of human RBPs and SARS-CoV-2 proteins that could alter the post-transcriptional regulatory mechanisms at several layers in the infected cells.

We show that the expression profiles of majority of the directly interacting RBPs were associated with gonadal tissues and immune cell types. Our study also highlights that several of the differentially expressed genes in SARS-CoV-2 infected cells were enriched for biological pathways such as immune response, cytokine mediated signaling, inflammatory response and metabolism associated genes that are indispensable for cell survival. Importantly, we found that among the differentially expressed genes, six RBP encoding genes contribute to functionally important pathways in the host cells, implying a potential impact of SARS-CoV-2 infection on post-transcriptional regulation. Further, our analysis demonstrates the abundance of skipped exonic and mutually exclusive exonic events in the SARS-CoV-2 infected cells, suggesting these alternative splicing events as a plausible cause for altered post-transcriptional regulation in human cells. Using motif enrichment analysis, we show that two key classes of post-transcriptional regulators, RBPs and miRs are likely to be titrated by SARS-CoV-2 genome that could result in systemic dysregulation of post-transcriptional networks in the infected cells. Currently, there are no effective anti-viral therapies available for COVID-19. Therefore, our analyses provide a roadmap to enhance the understanding of potential interactions of SARS-CoV-2 with key post-transcriptional regulators in the human genome.

## 4. Materials and methods

### 4.1. Dissection of SARS-CoV-2 proteins interacting with human RBPs

We obtained the high confidence mass spectrometry based SARS-CoV-2 viral protein to human protein interaction network established by Gordan et al, 2020 [23] in HEK293 cells. We dissected the human RBPs directly interacting with viral proteins and integrated with 1^st^ neighbor interacting RBPs (obtained from BioGRID [56]). We also extracted the protein abundance of these SARS-CoV-2 interacting RBPs across human tissues from Human Proteome Map [57]. The abundance of these proteins were hierarchically clustered and row normalized and represented as heatmap generated from Morpheus (https://software.broadinstitute.org/morpheus/).

### 4.2. Differential expression analysis of mock treated versus SARS-CoV-2 infected primary human lung epithelial cells

We downloaded the raw RNA sequencing data deposited in Gene Expression Omnibus (GEO)[65]. Specifically, we downloaded the paired end raw sequencing (FASTQ) files of mock treated and SARS-CoV-2 (USA-WA1/2020) infected primary human lung epithelial cells (in biological triplicates) using the Sequence Read Archive (SRA) Toolkit (fastq-dump command), from the GEO cohort GSE147507 [64]. The quality of the sequence reads were ensured using FASTX-Toolkit (http://hannonlab.cshl.edu/fastx_toolkit/) with a minimum of Phred quality score 20 for each sample. We processed the raw sequencing reads using the in-house NGS data processing pipeline as described previously[99, 100]. Briefly, we used Hierarchical Indexing for Spliced Alignment of Transcripts (HISAT) [101] for aligning the short reads from RNA-seq experiments onto human reference genome (hg38). SAM (Sequence Alignment/Map) files obtained from HISAT were post-processed using SAMtools (version 0.1.19) [102, 103] for converting SAM to BAM (Binary Alignment/Map) followed by sorting and indexing the output BAMs. The sorted BAM files were parsed using the python script provided by StringTie (version 1.2.1) [104] to obtain the count matrix of for gene levels across the samples. This count matrix was used to perform differential expression analysis between mock vs SARS-CoV-2 infected NHBE cells using DE-seq2 [105]. Statistically significant (at 5% fdr) differentially expressed genes were collected for downstream data analysis. Functional enrichment analysis of these genes was performed with p-value threshold < 10^−10^ using ClueGO [66] (a Cytoscape [106] plugin) andrepresented as bar-plot illustrating the significant pathways obtained from GO-term based functional grouping of differentially expressed genes.

### 4.3. Identification of alternative splicing events during SARS-CoV-2 infection

We used rMATS (replicate Multivariate Analysis of Transcript Splicing) [78] to identify differential alternative splicing (AS) events between the mock vs SARS-CoV-2 treated NHBE cells. rMATS used sorted BAM (Binary Alignment/Map) files, obtained from aligning the fastq files against hg38 reference genome using HISAT (as discussed above). It also uses a GTF file (gene transfer file format), downloaded from Ensembl (version 97) [107] for existing annotation of exons. Briefly, rMATS enabled the analysis of the inclusion/exclusion of target exons/introns contributing to different types of alternative splicing events, namely skipped exon (SE), alternative 5’ splice site (A5SS), alternative 3’ splice site (A3SS), mutually exclusive exons (MXE) and retained intron (RI), between pair of conditions and provides the difference in level of inclusion of an exon denoted by Percentage Splicing Index (*Ψ score*) (as described previously [99]). Genes exhibiting alternatively spliced events detected below 5% FDR threshold were documented in Table S2. Functional enrichment analysis of these genes was performed using ClueGO [66] and documented in Table S3.

### 4.4. Identification of potential binding blocks of RBPs in SARS-CoV-2 viral genome using motif enrichment analysis

We obtained the RBP-motifs from ATtRACT [83] and scanned across the SARS-CoV-2 viral genome using FIMO [82, 108] with default parameters. Resulting genomic locations for each RBP motif were documented in Table S4. For each binding motif, the scanned genomic location was normalized by considering the mid-point of genomic coordinate divided with the SARS-CoV-2 genome length. Also, the occurrence of each RBP motif binding across the viral genome was computed. Statistically significant (p-value < 1e-05) preferential binding profile of RBP motifs (sorted by frequency of binding and greater than 10 sites) across the SARS-CoV-2 viral genome (length normalized) identified using FIMO [82] was visualized in violin plot. Also, the protein abundance of corresponding RBPs were extracted from human proteome map[57] and represented as hierarchically clustered heatmap across the tissues.

### 4.5. Identification of potential binding blocks of microRNAs in SARS-CoV-2 viral genome using motif enrichment analysis

We obtained the miR-motifs from MEME [108] and scanned across the SARS-CoV-2 viral genome using FIMO[82, 108] with default parameters. Resulting genomic locations for each miR-motif were documented in Table S6. For each miR motif, the scanned genomic location was normalized by considering the mid-point of genomic coordinate divided with the SARS-CoV-2 genome length. Also, the occurrence of each miR-motif binding across the viral genome was computed. Statistically significant (p-value < 1e-05) preferential binding profile of miR-motifs (sorted by frequency of binding and greater than 15 sites) across the SARS-CoV-2 viral genome (length normalized) identified using FIMO was visualized in violin plot. Also, the expression profile of corresponding miRs were extracted from FANTOM5 project[109] and represented as hierarchically clustered heatmap across the tissues. To understand the generic biological function of these miRs, that could be altered by being titrated by SARS-CoV-2 genome in host cells, we downloaded the high confidence miR target genes (obtained from miRNet [110, 111]) and performed function annotation analysis. Resulting significant biological processes, obtained from gene ontology enrichment based functional grouping of these miR target genes were illustrated in barplot. Significant clustering (adj. p < 1e-10) of genes enriched in GO-biological processes were generated by ClueGO [66] analysis (a cytoscape [106] plugin).

## Acknowledgments

This work was supported by the National Institute of General Medical Sciences of the National Institutes of Health under Award Number R01GM123314 (SCJ), and COVID19 Rapid Response grant from IUPUI's Office of the Vice Chancellor for Research to SCJ. We also thank the lab members for their valuable suggestions and supporting dataset required for completion of this project.

## Authors’ contribution

RS, MS and SCJ conceived and designed the study. RS processed the publicly available dataset and implemented the bioinformatic tools. RS and SVD integrated the supplementary dataset required for downstream data analysis. RS, MS and SCJ interpreted the data and wrote the manuscript. All authors read and approved the final manuscript.

## References

1. de Wit, E.; van Doremalen, N.; Falzarano, D.; Munster, V. J. SARS and MERS: recent insights into emerging coronaviruses. Nat Rev Microbiol 2016, 14, (8), 523–34.

2. Meo, S. A.; Alhowikan, A. M.; Al-Khlaiwi, T.; Meo, I. M.; Halepoto, D. M.; Iqbal, M.; Usmani, M.; Hajjar, W.; Ahmed, N., Novel coronavirus 2019-nCoV: prevalence, biological and clinical characteristics comparison with SARS-CoV and MERS-CoV. Eur Rev Med Pharmacol Sci 2020, 24, (4), 2012–2019.

3. Zaki, A. M.; van Boheemen, S.; Bestebroer, T. M.; Osterhaus, A. D.; Fouchier, R. A. Isolation of a novel coronavirus from a man with pneumonia in Saudi Arabia. N Engl J Med 2012, 367, (19), 1814–20.

4. Cheng, V. C.; Lau, S. K.; Woo, P. C.; Yuen, K. Y. Severe acute respiratory syndrome coronavirus as an agent of emerging and reemerging infection. Clin Microbiol Rev 2007, 20, (4), 660–94.

5. Hajjar, S. A.; Memish, Z. A.; McIntosh, K. Middle East Respiratory Syndrome Coronavirus (MERS-CoV): a perpetual challenge. Ann Saudi Med 2013, 33, (5), 427–36.

6. Bolles, M.; Donaldson, E.; Baric, R., SARS-CoV and emergent coronaviruses: viral determinants of interspecies transmission. Curr Opin Virol 2011, 1, (6), 624–34.

7. Su, S.; Wong, G.; Shi, W.; Liu, J.; Lai, A. C. K.; Zhou, J.; Liu, W.; Bi, Y.; Gao, G. F., Epidemiology, Genetic Recombination, and Pathogenesis of Coronaviruses. Trends Microbiol 2016, 24, (6), 490–502.

8. Menachery, V. D.; Graham, R. L.; Baric, R. S., Jumping species-a mechanism for coronavirus persistence and survival. Curr Opin Virol 2017, 23, 1–7.

9. Zheng, J., SARS-CoV-2: an Emerging Coronavirus that Causes a Global Threat. Int J Biol Sci 2020, 16, (10), 1678–1685.

10. Xu, J.; Zhao, S.; Teng, T.; Abdalla, A. E.; Zhu, W.; Xie, L.; Wang, Y.; Guo, X. Systematic Comparison of Two Animal-to-Human Transmitted Human Coronaviruses: SARS-CoV-2 and SARS-CoV. Viruses 2020, 12, (2).

11. Gralinski, L. E.; Menachery, V. D. Return of the Coronavirus: 2019-nCoV. Viruses 2020, 12, (2).

12. Wang, H.; Li, X.; Li, T.; Zhang, S.; Wang, L.; Wu, X.; Liu, J. The genetic sequence, origin, and diagnosis of SARS-CoV-2. Eur J Clin Microbiol Infect Dis 2020, 39, (9), 1629–1635.

13. He, J.; Tao, H.; Yan, Y.; Huang, S. Y.; Xiao, Y. Molecular Mechanism of Evolution and Human Infection with SARS-CoV-2. Viruses 2020, 12, (4).

14. Trubsbach, A.; Dokert, B.; Schentke, K. U.; Jaross, W. [Lipid analysis of bile for determination of lithogeneity of bile and reproducibility of results with duodenal B bile]. Z Med Lab Diagn 1982, 23, (1), 30–4.

15. Kang, S.; Peng, W.; Zhu, Y.; Lu, S.; Zhou, M.; Lin, W.; Wu, W.; Huang, S.; Jiang, L.; Luo, X.; Deng, M. Recent progress in understanding 2019 novel coronavirus (SARS-CoV-2) associated with human respiratory disease: detection, mechanisms and treatment. Int J Antimicrob Agents 2020, 55, (5), 105950.

16. Malle, L. A map of SARS-CoV-2 and host cell interactions. Nat Rev Immunol 2020, 20, (6), 351.

17. Kim, D.; Lee, J. Y.; Yang, J. S.; Kim, J. W.; Kim, V. N.; Chang, H. The Architecture of SARS-CoV-2 Transcriptome. Cell 2020, 181, (4), 914–921 e10.

18. Khailany, R. A.; Safdar, M.; Ozaslan, M. Genomic characterization of a novel SARS-CoV-2. Gene Rep 2020, 19, 100682.

19. Elfiky, A. A., SARS-CoV-2 RNA dependent RNA polymerase (RdRp) targeting: an in silico perspective. J Biomol Struct Dyn 2020, 1–9.

20. Astuti, I.; Ysrafil, Severe Acute Respiratory Syndrome Coronavirus 2 (SARS-CoV-2): An overview of viral structure and host response. Diabetes Metab Syndr 2020, 14, (4), 407–412.

21. Guzzi, P. H.; Mercatelli, D.; Ceraolo, C.; Giorgi, F. M. Master Regulator Analysis of the SARS-CoV-2/Human Interactome. J Clin Med 2020, 9, (4).

22. Shereen, M. A.; Khan, S.; Kazmi, A.; Bashir, N.; Siddique, R., COVID-19 infection: Origin, transmission, and characteristics of human coronaviruses. J Adv Res 2020, 24, 91–98.

23. Gordon, D. E.; Jang, G. M.; Bouhaddou, M.; Xu, J.; Obernier, K.; White, K. M.; O’Meara, M. J.; Rezelj, V. V.; Guo, J. Z.; Swaney, D. L.; Tummino, T. A.; Huttenhain, R.; Kaake, R. M.; Richards, A. L.; Tutuncuoglu, B.; Foussard, H.; Batra, J.; Haas, K.; Modak, M.; Kim, M.; Haas, P.; Polacco, B. J.; Braberg, H.; Fabius, J. M.; Eckhardt, M.; Soucheray, M.; Bennett, M. J.; Cakir, M.; McGregor, M. J.; Li, Q.; Meyer, B.; Roesch, F.; Vallet, T.; Mac Kain, A.; Miorin, L.; Moreno, E.; Naing, Z. Z. C.; Zhou, Y.; Peng, S.; Shi, Y.; Zhang, Z.; Shen, W.; Kirby, I. T.; Melnyk, J. E.; Chorba, J. S.; Lou, K.; Dai, S. A.; Barrio-Hernandez, I.; Memon, D.; Hernandez-Armenta, C.; Lyu, J.; Mathy, C. J. P.; Perica, T.; Pilla, K. B.; Ganesan, S. J.; Saltzberg, D. J.; Rakesh, R.; Liu, X.; Rosenthal, S. B.; Calviello, L.; Venkataramanan, S.; Liboy-Lugo, J.; Lin, Y.; Huang, X. P.; Liu, Y.; Wankowicz, S. A.; Bohn, M.; Safari, M.; Ugur, F. S.; Koh, C.; Savar, N. S.; Tran, Q. D.; Shengjuler, D.; Fletcher, S. J.; O’Neal, M. C.; Cai, Y.; Chang, J. C. J.; Broadhurst, D. J.; Klippsten, S.; Sharp, P. P.; Wenzell, N. A.; Kuzuoglu-Ozturk, D.; Wang, H. Y.; Trenker, R.; Young, J. M.; Cavero, D. A.; Hiatt, J.; Roth, T. L.; Rathore, U.; Subramanian, A.; Noack, J.; Hubert, M.; Stroud, R. M.; Frankel, A. D.; Rosenberg, O. S.; Verba, K. A.; Agard, D. A.; Ott, M.; Emerman, M.; Jura, N.; von Zastrow, M.; Verdin, E.; Ashworth, A.; Schwartz, O.; d’Enfert, C.; Mukherjee, S.; Jacobson, M.; Malik, H. S.; Fujimori, D. G.; Ideker, T.; Craik, C. S.; Floor, S. N.; Fraser, J. S.; Gross, J. D.; Sali, A.; Roth, B. L.; Ruggero, D.; Taunton, J.; Kortemme, T.; Beltrao, P.; Vignuzzi, M.; Garcia-Sastre, A.; Shokat, K. M.; Shoichet, B. K.; Krogan, N. J., A SARS-CoV-2 protein interaction map reveals targets for drug repurposing. Nature 2020, 583, (7816), 459–468.

24. Hentze, M. W.; Castello, A.; Schwarzl, T.; Preiss, T. A brave new world of RNA-binding proteins. Nat Rev Mol Cell Biol 2018, 19, (5), 327–341.

25. Dassi, E. Handshakes and Fights: The Regulatory Interplay of RNA-Binding Proteins. Front Mol Biosci 2017, 4, 67.

26. Corley, M.; Burns, M. C.; Yeo, G. W., How RNA-Binding Proteins Interact with RNA: Molecules and Mechanisms. Mol Cell 2020, 78, (1), 9–29.

27. O’Brien, J.; Hayder, H.; Zayed, Y.; Peng, C. Overview of MicroRNA Biogenesis, Mechanisms of Actions, and Circulation. Front Endocrinol (Lausanne) 2018, 9, 402.

28. Hammond, S. M. An overview of microRNAs. Adv Drug Deliv Rev 2015, 87, 3–14.

29. Glisovic, T.; Bachorik, J. L.; Yong, J.; Dreyfuss, G., RNA-binding proteins and post-transcriptional gene regulation. FEBS Lett 2008, 582, (14), 1977–86.

30. Zanzoni, A.; Spinelli, L.; Ribeiro, D. M.; Tartaglia, G. G.; Brun, C., Post-transcriptional regulatory patterns revealed by protein-RNA interactions. Sci Rep 2019, 9, (1), 4302.

31. Chen, P. Y.; Meister, G., microRNA-guided posttranscriptional gene regulation. Biol Chem 2005, 386, (12), 1205–18.

32. Brinegar, A. E.; Cooper, T. A. Roles for RNA-binding proteins in development and disease. Brain Res 2016, 1647, 1–8.

33. Lukong, K. E.; Chang, K. W.; Khandjian, E. W.; Richard, S., RNA-binding proteins in human genetic disease. Trends Genet 2008, 24, (8), 416–25.

34. Li, Z.; Nagy, P. D. Diverse roles of host RNA binding proteins in RNA virus replication. RNA Biol 2011, 8, (2), 305–15.

35. Garcia-Moreno, M.; Noerenberg, M.; Ni, S.; Jarvelin, A. I.; Gonzalez-Almela, E.; Lenz, C. E.; Bach-Pages, M.; Cox, V.; Avolio, R.; Davis, T.; Hester, S.; Sohier, T. J. M.; Li, B.; Heikel, G.; Michlewski, G.; Sanz, M. A.; Carrasco, L.; Ricci, E. P.; Pelechano, V.; Davis, I.; Fischer, B.; Mohammed, S.; Castello, A., System-wide Profiling of RNA-Binding Proteins Uncovers Key Regulators of Virus Infection. Mol Cell 2019, 74, (1), 196–211 e11.

36. Ardekani, A. M.; Naeini, M. M. The Role of MicroRNAs in Human Diseases. Avicenna J Med Biotechnol 2010, 2, (4), 161–79.

37. Drury, R. E.; O’Connor, D.; Pollard, A. J. The Clinical Application of MicroRNAs in Infectious Disease. Front Immunol 2017, 8, 1182.

38. Girardi, E.; Lopez, P.; Pfeffer, S. On the Importance of Host MicroRNAs During Viral Infection. Front Genet 2018, 9, 439.

39. Srinivasan, S.; Cui, H.; Gao, Z.; Liu, M.; Lu, S.; Mkandawire, W.; Narykov, O.; Sun, M.; Korkin, D. Structural Genomics of SARS-CoV-2 Indicates Evolutionary Conserved Functional Regions of Viral Proteins. Viruses 2020, 12, (4).

40. Maranon, D. G.; Anderson, J. R.; Maranon, A. G.; Wilusz, J. The interface between coronaviruses and host cell RNA biology: Novel potential insights for future therapeutic intervention. Wiley Interdiscip Rev RNA 2020, 11, (5), e1614.

41. Mukherjee, M.; Goswami, S. Global cataloguing of variations in untranslated regions of viral genome and prediction of key host RNA binding protein-microRNA interactions modulating genome stability in SARS-CoV-2. PLoS One 2020, 15, (8), e0237559.

42. Luo, H.; Chen, Q.; Chen, J.; Chen, K.; Shen, X.; Jiang, H. The nucleocapsid protein of SARS coronavirus has a high binding affinity to the human cellular heterogeneous nuclear ribonucleoprotein A1. FEBS Lett 2005, 579, (12), 2623–8.

43. Sola, I.; Galan, C.; Mateos-Gomez, P. A.; Palacio, L.; Zuniga, S.; Cruz, J. L.; Almazan, F.; Enjuanes, L. The polypyrimidine tract-binding protein affects coronavirus RNA accumulation levels and relocalizes viral RNAs to novel cytoplasmic domains different from replication-transcription sites. J Virol 2011, 85, (10), 5136–49.

44. Sola, I.; Mateos-Gomez, P. A.; Almazan, F.; Zuniga, S.; Enjuanes, L., RNA-RNA and RNA-protein interactions in coronavirus replication and transcription. RNA Biol 2011, 8, (2), 237–48.

45. Mallick, B.; Ghosh, Z.; Chakrabarti, J. MicroRNome analysis unravels the molecular basis of SARS infection in bronchoalveolar stem cells. PLoS One 2009, 4, (11), e7837.

46. Peng, X.; Gralinski, L.; Ferris, M. T.; Frieman, M. B.; Thomas, M. J.; Proll, S.; Korth, M. J.; Tisoncik, J. R.; Heise, M.; Luo, S.; Schroth, G. P.; Tumpey, T. M.; Li, C.; Kawaoka, Y.; Baric, R. S.; Katze, M. G. Integrative deep sequencing of the mouse lung transcriptome reveals differential expression of diverse classes of small RNAs in response to respiratory virus infection. mBio 2011, 2, (6).

47. Wu, C.; Liu, Y.; Yang, Y.; Zhang, P.; Zhong, W.; Wang, Y.; Wang, Q.; Xu, Y.; Li, M.; Li, X.; Zheng, M.; Chen, L.; Li, H. Analysis of therapeutic targets for SARS-CoV-2 and discovery of potential drugs by computational methods. Acta Pharm Sin B 2020, 10, (5), 766–788.

48. Gurwitz, D. Angiotensin receptor blockers as tentative SARS-CoV-2 therapeutics. Drug Dev Res 2020, 81, (5), 537–540.

49. Shi, S. T.; Lai, M. M. Viral and cellular proteins involved in coronavirus replication. Curr Top Microbiol Immunol 2005, 287, 95–131.

50. Spagnolo, J. F.; Hogue, B. G. Host protein interactions with the 3' end of bovine coronavirus RNA and the requirement of the poly(A) tail for coronavirus defective genome replication. J Virol 2000, 74, (11), 5053–65.

51. Yu, W.; Leibowitz, J. L. Specific binding of host cellular proteins to multiple sites within the 3’ end of mouse hepatitis virus genomic RNA. J Virol 1995, 69, (4), 2016–23.

52. Tutuncuoglu, B.; Cakir, M.; Batra, J.; Bouhaddou, M.; Eckhardt, M.; Gordon, D. E.; Krogan, N. J. The Landscape of Human Cancer Proteins Targeted by SARS-CoV-2. Cancer Discov 2020, 10, (7), 916–921.

53. Nyathi, Y.; Wilkinson, B. M.; Pool, M. R., Co-translational targeting and translocation of proteins to the endoplasmic reticulum. Biochimica et biophysica acta 2013, 1833, (11), 2392–402.

54. Saraogi, I.; Shan, S. O. Molecular mechanism of co-translational protein targeting by the signal recognition particle. Traffic 2011, 12, (5), 535–42.

55. Hashemikhabir, S.; Neelamraju, Y.; Janga, S. C. Database of RNA binding protein expression and disease dynamics (READ DB). Database : the journal of biological databases and curation 2015, 2015, bav072.

56. Chatr-Aryamontri, A.; Breitkreutz, B. J.; Oughtred, R.; Boucher, L.; Heinicke, S.; Chen, D.; Stark, C.; Breitkreutz, A.; Kolas, N.; O’Donnell, L.; Reguly, T.; Nixon, J.; Ramage, L.; Winter, A.; Sellam, A.; Chang, C.; Hirschman, J.; Theesfeld, C.; Rust, J.; Livstone, M. S.; Dolinski, K.; Tyers, M. The BioGRID interaction database: 2015 update. Nucleic Acids Res 2015, 43, (Database issue), D470–8.

57. Kim, M. S.; Pinto, S. M.; Getnet, D.; Nirujogi, R. S.; Manda, S. S.; Chaerkady, R.; Madugundu, K.; Kelkar, D. S.; Isserlin, R.; Jain, S.; Thomas, J. K.; Muthusamy, B.; Leal-Rojas, P.; Kumar, P.; Sahasrabuddhe, N. A.; Balakrishnan, L.; Advani, J.; George, B.; Renuse, S.; Selvan, L. D.; Patil, A. H.; Nanjappa, V.; Radhakrishnan, A.; Prasad, S.; Subbannayya, T.; Raju, R.; Kumar, M.; Sreenivasamurthy, S. K.; Marimuthu, A.; Sathe, G. J.; Chavan, S.; Datta, K. K.; Subbannayya, Y.; Sahu, A.; Yelamanchi, S. D.; Jayaram, S.; Rajagopalan, P.; Sharma, J.; Murthy, K. R.; Syed, N.; Goel, R.; Khan, A. A.; Ahmad, S.; Dey, G.; Mudgal, K.; Chatterjee, A.; Huang, T. C.; Zhong, J.; Wu, X.; Shaw, P. G.; Freed, D.; Zahari, M. S.; Mukherjee, K. K.; Shankar, S.; Mahadevan, A.; Lam, H.; Mitchell, C. J.; Shankar, S. K.; Satishchandra, P.; Schroeder, J. T.; Sirdeshmukh, R.; Maitra, A.; Leach, S. D.; Drake, C. G.; Halushka, M. K.; Prasad, T. S.; Hruban, R. H.; Kerr, C. L.; Bader, G. D.; Iacobuzio-Donahue, C. A.; Gowda, H.; Pandey, A. A draft map of the human proteome. Nature 2014, 509, (7502), 575–81.

58. Wang, S.; Zhou, X.; Zhang, T.; Wang, Z. The need for urogenital tract monitoring in COVID-19. Nat Rev Urol 2020, 17, (6), 314–315.

59. Ma, L.; Xie, W.; Li, D.; Shi, L.; Mao, Y.; Xiong, Y.; Zhang, Y.; Zhang, M. Effect of SARS-CoV-2 infection upon male gonadal function: A single center-based study. medRxiv 2020.

60. Li, G.; Fan, Y.; Lai, Y.; Han, T.; Li, Z.; Zhou, P.; Pan, P.; Wang, W.; Hu, D.; Liu, X.; Zhang, Q.; Wu, J. Coronavirus infections and immune responses. J Med Virol 2020, 92, (4), 424–432.

61. Wang, X.; Xu, W.; Hu, G.; Xia, S.; Sun, Z.; Liu, Z.; Xie, Y.; Zhang, R.; Jiang, S.; Lu, L., RETRACTED ARTICLE: SARS-CoV-2 infects T lymphocytes through its spike protein-mediated membrane fusion. Cell Mol Immunol 2020.

62. Zheng, M.; Gao, Y.; Wang, G.; Song, G.; Liu, S.; Sun, D.; Xu, Y.; Tian, Z. Functional exhaustion of antiviral lymphocytes in COVID-19 patients. Cell Mol Immunol 2020, 17, (5), 533–535.

63. Braun, J.; Loyal, L.; Frentsch, M.; Wendisch, D.; Georg, P.; Kurth, F.; Hippenstiel, S.; Dingeldey, M.; Kruse, B.; Fauchere, F.; Baysal, E.; Mangold, M.; Henze, L.; Lauster, R.; Mall, M. A.; Beyer, K.; Rohmel, J.; Voigt, S.; Schmitz, J.; Miltenyi, S.; Demuth, I.; Muller, M. A.; Hocke, A.; Witzenrath, M.; Suttorp, N.; Kern, F.; Reimer, U.; Wenschuh, H.; Drosten, C.; Corman, V. M.; Giesecke-Thiel, C.; Sander, L. E.; Thiel, A., SARS-CoV-2-reactive T cells in healthy donors and patients with COVID-19. Nature 2020.

64. Blanco-Melo, D.; Nilsson-Payant, B. E.; Liu, W. C.; Uhl, S.; Hoagland, D.; Moller, R.; Jordan, T. X.; Oishi, K.; Panis, M.; Sachs, D.; Wang, T. T.; Schwartz, R. E.; Lim, J. K.; Albrecht, R. A.; tenOever, B. R. Imbalanced Host Response to SARS-CoV-2 Drives Development of COVID-19. Cell 2020, 181, (5), 1036–1045 e9.

65. Barrett, T.; Wilhite, S. E.; Ledoux, P.; Evangelista, C.; Kim, I. F.; Tomashevsky, M.; Marshall, K. A.; Phillippy, K. H.; Sherman, P. M.; Holko, M.; Yefanov, A.; Lee, H.; Zhang, N.; Robertson, C. L.; Serova, N.; Davis, S.; Soboleva, A., NCBI GEO: archive for functional genomics data sets--update. Nucleic Acids Res 2013, 41, (Database issue), D991–5.

66. Bindea, G.; Mlecnik, B.; Hackl, H.; Charoentong, P.; Tosolini, M.; Kirilovsky, A.; Fridman, W. H.; Pages, F.; Trajanoski, Z.; Galon, J., ClueGO: a Cytoscape plug-in to decipher functionally grouped gene ontology and pathway annotation networks. Bioinformatics 2009, 25, (8), 1091–3.

67. Xu, Z.; Shi, L.; Wang, Y.; Zhang, J.; Huang, L.; Zhang, C.; Liu, S.; Zhao, P.; Liu, H.; Zhu, L.; Tai, Y.; Bai, C.; Gao, T.; Song, J.; Xia, P.; Dong, J.; Zhao, J.; Wang, F. S. Pathological findings of COVID-19 associated with acute respiratory distress syndrome. Lancet Respir Med 2020, 8, (4), 420–422.

68. Zhang, C.; Wu, Z.; Li, J. W.; Zhao, H.; Wang, G. Q. Cytokine release syndrome in severe COVID-19: interleukin-6 receptor antagonist tocilizumab may be the key to reduce mortality. Int J Antimicrob Agents 2020, 55, (5), 105954.

69. Wu, D.; Yang, X. O., TH17 responses in cytokine storm of COVID-19: An emerging target of JAK2 inhibitor Fedratinib. J Microbiol Immunol Infect 2020, 53, (3), 368–370.

70. Kudva, A.; Scheller, E. V.; Robinson, K. M.; Crowe, C. R.; Choi, S. M.; Slight, S. R.; Khader, S. A.; Dubin, P. J.; Enelow, R. I.; Kolls, J. K.; Alcorn, J. F. Influenza A inhibits Th17-mediated host defense against bacterial pneumonia in mice. J Immunol 2011, 186, (3), 1666–1674.

71. Pacha, O.; Sallman, M. A.; Evans, S. E., COVID-19: a case for inhibiting IL-17? Nat Rev Immunol 2020, 20, (6), 345–346.

72. Chauhan, K.; Kalam, H.; Dutt, R.; Kumar, D., RNA Splicing: A New Paradigm in Host-Pathogen Interactions. Journal of molecular biology 2019, 431, (8), 1565–1575.

73. Ashraf, U.; Benoit-Pilven, C.; Lacroix, V.; Navratil, V.; Naffakh, N. Advances in Analyzing Virus-Induced Alterations of Host Cell Splicing. Trends Microbiol 2019, 27, (3), 268–281.

74. Boudreault, S.; Roy, P.; Lemay, G.; Bisaillon, M. Viral modulation of cellular RNA alternative splicing: A new key player in virus-host interactions? Wiley Interdiscip Rev RNA 2019, 10, (5), e1543.

75. Bojkova, D.; Klann, K.; Koch, B.; Widera, M.; Krause, D.; Ciesek, S.; Cinatl, J.; Münch, C., SARS-CoV-2 infected host cell proteomics reveal potential therapy targets. 2020.

76. Lim, Y. X.; Ng, Y. L.; Tam, J. P.; Liu, D. X., Human Coronaviruses: A Review of Virus-Host Interactions. Diseases 2016, 4, (3).

77. DeDiego, M. L.; Nieto-Torres, J. L.; Jimenez-Guardeno, J. M.; Regla-Nava, J. A.; Alvarez, E.; Oliveros, J. C.; Zhao, J.; Fett, C.; Perlman, S.; Enjuanes, L. Severe acute respiratory syndrome coronavirus envelope protein regulates cell stress response and apoptosis. PLoS Pathog 2011, 7, (10), e1002315.

78. Shen, S.; Park, J. W.; Lu, Z. X.; Lin, L.; Henry, M. D.; Wu, Y. N.; Zhou, Q.; Xing, Y., rMATS: robust and flexible detection of differential alternative splicing from replicate RNA-Seq data. Proceedings of the National Academy of Sciences of the United States of America 2014, 111, (51), E5593–601.

79. Forster, P.; Forster, L.; Renfrew, C.; Forster, M. Phylogenetic network analysis of SARS-CoV-2 genomes. Proceedings of the National Academy of Sciences of the United States of America 2020, 117, (17), 9241–9243.

80. Zhu, J.; Gopinath, K.; Murali, A.; Yi, G.; Hayward, S. D.; Zhu, H.; Kao, C., RNA-binding proteins that inhibit RNA virus infection. Proceedings of the National Academy of Sciences of the United States of America 2007, 104, (9), 3129–34.

81. Garcia-Moreno, M.; Jarvelin, A. I.; Castello, A., Unconventional RNA-binding proteins step into the virus-host battlefront. Wiley Interdiscip Rev RNA 2018, 9, (6), e1498.

82. Grant, C. E.; Bailey, T. L.; Noble, W. S., FIMO: scanning for occurrences of a given motif. Bioinformatics 2011, 27, (7), 1017–8.

83. Giudice, G.; Sanchez-Cabo, F.; Torroja, C.; Lara-Pezzi, E., ATtRACT-a database of RNA-binding proteins and associated motifs. Database : the journal of biological databases and curation 2016, 2016.

84. Castello, A.; Fischer, B.; Eichelbaum, K.; Horos, R.; Beckmann, B. M.; Strein, C.; Davey, N. E.; Humphreys, D. T.; Preiss, T.; Steinmetz, L. M.; Krijgsveld, J.; Hentze, M. W. Insights into RNA biology from an atlas of mammalian mRNA-binding proteins. Cell 2012, 149, (6), 1393–406.

85. Bartas, M.; Brazda, V.; Bohalova, N.; Cantara, A.; Volna, A.; Stachurova, T.; Malachova, K.; Jagelska, E. B.; Porubiakova, O.; Cerven, J.; Pecinka, P., In-Depth Bioinformatic Analyses of Nidovirales Including Human SARS-CoV-2, SARS-CoV, MERS-CoV Viruses Suggest Important Roles of Non-canonical Nucleic Acid Structures in Their Lifecycles. Front Microbiol 2020, 11, (1583), 1583.

86. Sapra, A. K.; Anko, M. L.; Grishina, I.; Lorenz, M.; Pabis, M.; Poser, I.; Rollins, J.; Weiland, E. M.; Neugebauer, K. M. SR protein family members display diverse activities in the formation of nascent and mature mRNPs in vivo. Mol Cell 2009, 34, (2), 179–90.

87. Cazalla, D.; Zhu, J.; Manche, L.; Huber, E.; Krainer, A. R.; Caceres, J. F. Nuclear export and retention signals in the RS domain of SR proteins. Molecular and cellular biology 2002, 22, (19), 6871–82.

88. Sanford, J. R.; Gray, N. K.; Beckmann, K.; Caceres, J. F. A novel role for shuttling SR proteins in mRNA translation. Genes Dev 2004, 18, (7), 755–68.

89. Michlewski, G.; Sanford, J. R.; Caceres, J. F. The splicing factor SF2/ASF regulates translation initiation by enhancing phosphorylation of 4E-BP1. Mol Cell 2008, 30, (2), 179–89.

90. Botti, V.; McNicoll, F.; Steiner, M. C.; Richter, F. M.; Solovyeva, A.; Wegener, M.; Schwich, O. D.; Poser, I.; Zarnack, K.; Wittig, I.; Neugebauer, K. M.; Muller-McNicoll, M. Cellular differentiation state modulates the mRNA export activity of SR proteins. J Cell Biol 2017, 216, (7), 1993–2009.

91. Caceres, J. F.; Screaton, G. R.; Krainer, A. R. A specific subset of SR proteins shuttles continuously between the nucleus and the cytoplasm. Genes Dev 1998, 12, (1), 55–66.

92. Dickson, A. M.; Wilusz, J. Strategies for viral RNA stability: live long and prosper. Trends Genet 2011, 27, (7), 286–93.

93. Bruscella, P.; Bottini, S.; Baudesson, C.; Pawlotsky, J. M.; Feray, C.; Trabucchi, M. Viruses and miRNAs: More Friends than Foes. Front Microbiol 2017, 8, 824.

94. Roberts, A. P.; Lewis, A. P.; Jopling, C. L. The role of microRNAs in viral infection. Prog Mol Biol Transl Sci 2011, 102, 101–39.

95. Sacar Demirci, M. D.; Adan, A. Computational analysis of microRNA-mediated interactions in SARS-CoV-2 infection. PeerJ 2020, 8, e9369.

96. Khan, M. A.; Sany, M. R. U.; Islam, M. S.; Islam, A. Epigenetic Regulator miRNA Pattern Differences Among SARS-CoV, SARS-CoV-2, and SARS-CoV-2 World-Wide Isolates Delineated the Mystery Behind the Epic Pathogenicity and Distinct Clinical Characteristics of Pandemic COVID-19. Front Genet 2020, 11, 765.

97. Xu, X.; Chen, P.; Wang, J.; Feng, J.; Zhou, H.; Li, X.; Zhong, W.; Hao, P. Evolution of the novel coronavirus from the ongoing Wuhan outbreak and modeling of its spike protein for risk of human transmission. Sci China Life Sci 2020, 63, (3), 457–460.

98. Graham, R. L.; Sparks, J. S.; Eckerle, L. D.; Sims, A. C.; Denison, M. R. SARS coronavirus replicase proteins in pathogenesis. Virus Res 2008, 133, (1), 88–100.

99. Srivastava, R.; Budak, G.; Dash, S.; Lachke, S. A.; Janga, S. C. Transcriptome analysis of developing lens reveals abundance of novel transcripts and extensive splicing alterations. Sci Rep 2017, 7, (1), 11572.

100. Budak, G.; Dash, S.; Srivastava, R.; Lachke, S. A.; Janga, S. C., Express: A database of transcriptome profiles encompassing known and novel transcripts across multiple development stages in eye tissues. Experimental eye research 2018, 168, 57–68.

101. Kim, D.; Langmead, B.; Salzberg, S. L., HISAT: a fast spliced aligner with low memory requirements. Nat Methods 2015, 12, (4), 357–60.

102. Li, H.; Handsaker, B.; Wysoker, A.; Fennell, T.; Ruan, J.; Homer, N.; Marth, G.; Abecasis, G.; Durbin, R.; Genome Project Data Processing, S. The Sequence Alignment/Map format and SAMtools. Bioinformatics 2009, 25, (16), 2078–9.

103. Wheeler, D. L.; Barrett, T.; Benson, D. A.; Bryant, S. H.; Canese, K.; Chetvernin, V.; Church, D. M.; Dicuccio, M.; Edgar, R.; Federhen, S.; Feolo, M.; Geer, L. Y.; Helmberg, W.; Kapustin, Y.; Khovayko, O.; Landsman, D.; Lipman, D. J.; Madden, T. L.; Maglott, D. R.; Miller, V.; Ostell, J.; Pruitt, K. D.; Schuler, G. D.; Shumway, M.; Sequeira, E.; Sherry, S. T.; Sirotkin, K.; Souvorov, A.; Starchenko, G.; Tatusov, R. L.; Tatusova, T. A.; Wagner, L.; Yaschenko, E. Database resources of the National Center for Biotechnology Information. Nucleic Acids Res 2008, 36, (Database issue), D13–21.

104. Pertea, M.; Pertea, G. M.; Antonescu, C. M.; Chang, T. C.; Mendell, J. T.; Salzberg, S. L. StringTie enables improved reconstruction of a transcriptome from RNA-seq reads. Nat Biotechnol 2015, 33, (3), 290–5.

105. Love, M. I.; Huber, W.; Anders, S. Moderated estimation of fold change and dispersion for RNA-seq data with DESeq2. Genome biology 2014, 15, (12), 550.

106. Smoot, M. E.; Ono, K.; Ruscheinski, J.; Wang, P. L.; Ideker, T., Cytoscape 2.8: new features for data integration and network visualization. Bioinformatics 2011, 27, (3), 431–2.

107. Cunningham, F.; Amode, M. R.; Barrell, D.; Beal, K.; Billis, K.; Brent, S.; Carvalho-Silva, D.; Clapham, P.; Coates, G.; Fitzgerald, S.; Gil, L.; Giron, C. G.; Gordon, L.; Hourlier, T.; Hunt, S. E.; Janacek, S. H.; Johnson, N.; Juettemann, T.; Kahari, A. K.; Keenan, S.; Martin, F. J.; Maurel, T.; McLaren, W.; Murphy, D. N.; Nag, R.; Overduin, B.; Parker, A.; Patricio, M.; Perry, E.; Pignatelli, M.; Riat, H. S.; Sheppard, D.; Taylor, K.; Thormann, A.; Vullo, A.; Wilder, S. P.; Zadissa, A.; Aken, B. L.; Birney, E.; Harrow, J.; Kinsella, R.; Muffato, M.; Ruffier, M.; Searle, S. M.; Spudich, G.; Trevanion, S. J.; Yates, A.; Zerbino, D. R.; Flicek, P., Ensembl 2015. Nucleic Acids Res 2015, 43, (Database issue), D662–9.

108. Bailey, T. L.; Johnson, J.; Grant, C. E.; Noble, W. S. The MEME Suite. Nucleic Acids Res 2015, 43, (W1), W39–49.

109. de Rie, D.; Abugessaisa, I.; Alam, T.; Arner, E.; Arner, P.; Ashoor, H.; Astrom, G.; Babina, M.; Bertin, N.; Burroughs, A. M.; Carlisle, A. J.; Daub, C. O.; Detmar, M.; Deviatiiarov, R.; Fort, A.; Gebhard, C.; Goldowitz, D.; Guhl, S.; Ha, T. J.; Harshbarger, J.; Hasegawa, A.; Hashimoto, K.; Herlyn, M.; Heutink, P.; Hitchens, K. J.; Hon, C. C.; Huang, E.; Ishizu, Y.; Kai, C.; Kasukawa, T.; Klinken, P.; Lassmann, T.; Lecellier, C. H.; Lee, W.; Lizio, M.; Makeev, V.; Mathelier, A.; Medvedeva, Y. A.; Mejhert, N.; Mungall, C. J.; Noma, S.; Ohshima, M.; Okada-Hatakeyama, M.; Persson, H.; Rizzu, P.; Roudnicky, F.; Saetrom, P.; Sato, H.; Severin, J.; Shin, J. W.; Swoboda, R. K.; Tarui, H.; Toyoda, H.; Vitting-Seerup, K.; Winteringham, L.; Yamaguchi, Y.; Yasuzawa, K.; Yoneda, M.; Yumoto, N.; Zabierowski, S.; Zhang, P. G.; Wells, C. A.; Summers, K. M.; Kawaji, H.; Sandelin, A.; Rehli, M.; Consortium, F.; Hayashizaki, Y.; Carninci, P.; Forrest, A. R. R.; de Hoon, M. J. L., An integrated expression atlas of miRNAs and their promoters in human and mouse. Nat Biotechnol 2017, 35, (9), 872–878.

110. Fan, Y.; Siklenka, K.; Arora, S. K.; Ribeiro, P.; Kimmins, S.; Xia, J., miRNet - dissecting miRNA-target interactions and functional associations through network-based visual analysis. Nucleic Acids Res 2016, 44, (W1), W135–41.

111. Fan, Y.; Xia, J., miRNet-Functional Analysis and Visual Exploration of miRNA-Target Interactions in a Network Context. Methods Mol Biol 2018, 1819, 215–233.

